# Integrating environmental and ecological monitoring with seaweed farming

**DOI:** 10.1101/2024.02.15.580450

**Authors:** Christian Berger, Polina Papazova, Beth Marshall, Fergus Evans, Craig Williams

## Abstract

We explore the biodiversity impacts of seaweed and shellfish farms in Pembrokeshire, UK, with a focus on using monitoring methods that are affordable and can be integrated into existing seaweed farming operations. Monitoring methods used include Baited Remote Underwater Video Systems (BRUVs), Passive Acoustic Monitoring (PAM) and visual surveys of cultivation lines and natural settlement on farm infrastructure. BRUVs detected 13 motile fauna taxa, influenced mainly by site conditions, while PAM observed distinct patterns in dolphin and porpoise activity that changes between seasons. Visual surveys revealed colonisation of algal, hydroid, and bryozoan species on both the surface and seabed infrastructure, indicating that the farming structures serve as stable substrates for biodiversity accumulation. Apart from the natural settlement on infrastructure, our data does not show a conclusive link between biodiversity and farming due to the highly dynamic environment, small scale of operations, and relatively short monitoring timeframe. Despite these limitations, the data sets a crucial baseline for future studies and showed no negative impacts even as farming activities intensified over the monitoring period. The study aligns with what’s feasible for time-constrained farm operators and forms a foundation for routine, integrated environmental and biodiversity monitoring.

## 1. Introduction

Seaweed farming is an emergent and rapidly growing sector of aquaculture, contributing to a significant percentage of global aquaculture biomass and displaying a steady annual growth rate of around 8% [1]. The cultivation of seaweed is economically promising and has well-documented environmental benefits such as carbon sequestration, nutrient cycling, and habitat provision [2], [3], [4], [5]. Research on understanding these ecosystem services is beginning to gather pace and a consensus on the roles of carbon and nutrient cycling are beginning to emerge [6], [7]. However, the impacts on marine biodiversity remains a very complex picture, and in the context of the northeast Atlantic region, there are a limited number of comprehensive studies with inconclusive findings that show highly variable observations across different taxa [3], [8].

The urgency to prioritise research in this area is underscored by accumulating evidence of biodiversity loss of vulnerable marine habitats and the emergence of more proactive policies on biodiversity, such as the UK’s Marine Net Gain policy [9], [10]. Without sufficient data on seaweed farming, we risk missing out on its potential benefits, as such projects may go unrecognised and underfunded. A key challenge in this area of research is the lack of high-quality, large-scale, and long-term environmental and biodiversity data. The existing literature often faces methodological limitations such as the absence of baseline reference data and the inadequacy of streamlined workflows for monitoring [11]. These gaps in knowledge and methodology pose significant barriers to the development of evidence-based policies and sustainable aquaculture practices [12]. Recognising the complexities and gaps in existing literature, along with the current reliance on expensive scientific research conducted by experts, our study seeks to establish a practical, multi-method approach to make biodiversity monitoring an integral part of seaweed farming operations.

With the aim of making environmental and biodiversity monitoring more accessible and routine, we assessed seaweed using direct observations, Baited Remote Underwater Video (BRUV) and Passive Acoustic Monitoring (PAM). These methods were selected as they are cost-effective, easy to integrate into existing seaweed farming operations, and capable of capturing a wide range of species in terms of size and mobility. Moreover, they are well-suited for our study area’s dynamic, sediment-poor marine conditions, which make traditional methods like sediment microscopy challenging. Whilst we also considered environmental DNA (eDNA) sampling as a method to identifying a broader range of organisms, the absence of sediment at the study sites made it difficult to collect dependable eDNA samples. Water samples for eDNA analysis were also not included in our study due to the strong currents at the sites, limiting the ability to capture DNA from sources local to the seaweed farm. Nevertheless, as seaweed farms grow into offshore areas with more sediment and slower currents, eDNA could become an important tool for future monitoring.

**Fig. 1.**
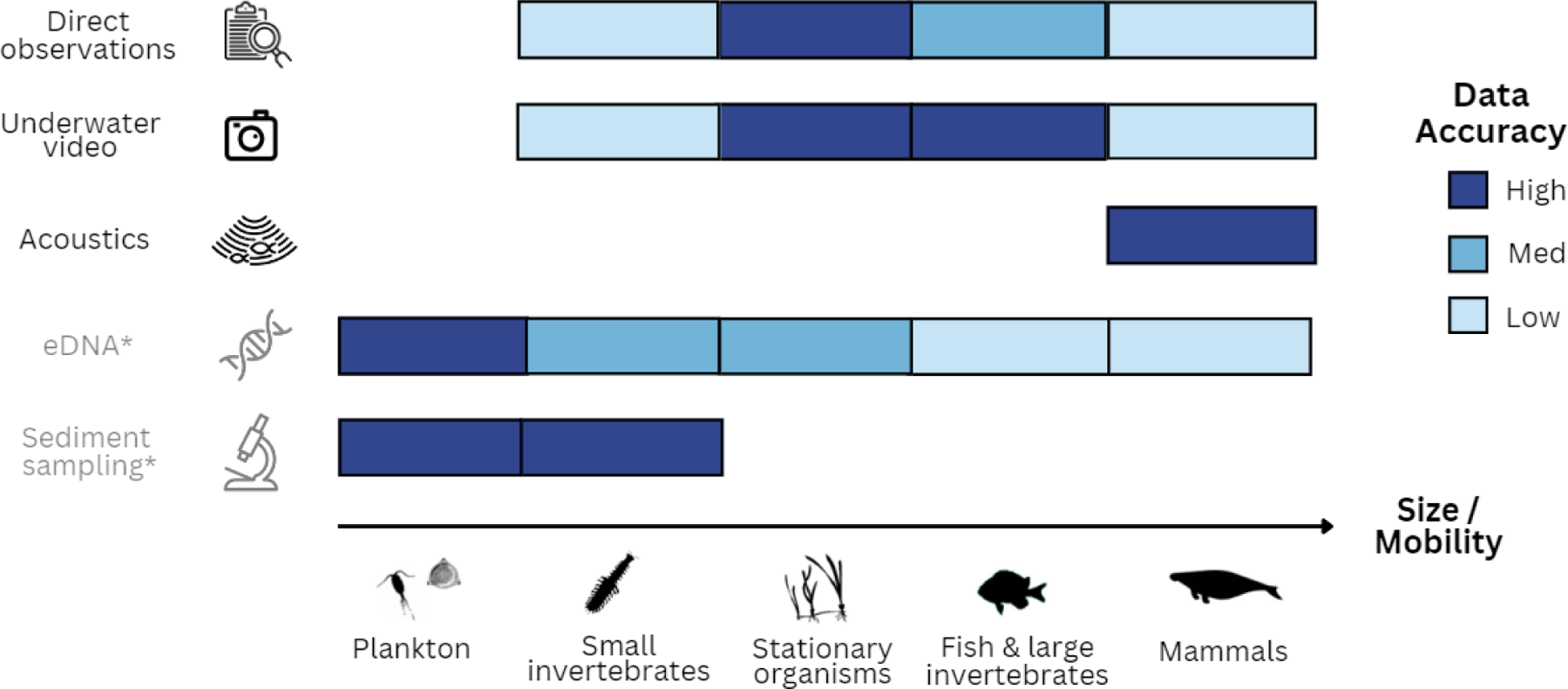
Biodiversity monitoring methods considered in this study, including their accuracy in capturing biodiversity at different size and mobility scales. The coloured bars represent the accuracy at which each method can detect and measure the corresponding organisms. *eDNA and sediment sampling were not carried as part of this study.

An overview of the biodiversity monitoring methods employed in this study is provided in Fig. 1, describing the accuracy of data captured for organisms of varying size and mobility across each method considered. In the current study, selection of observation methods was based on their ability to yield high-accuracy data. Accordingly, visual surveys were used to monitor stationary organisms, video techniques were employed for observing stationary organisms as well as fish and large invertebrates, and acoustic monitoring was reserved for the study of marine mammals. Additionally, we collected data on physical environmental parameters to provide context for our biodiversity observations.

### 1.1 Direct observations

Organisms that foul seaweeds and their associated farming infrastructure are typically considered as a pest, reducing crop biomass and quality [13], [14], [15]. Nevertheless, specific epibionts like bivalves and bryozoans have the potential to positively impact the local marine environment. They can improve water quality through biofiltration and serve as a food source for benthic organisms when they dislodge from the seaweed farm canopy and settle on the ocean floor [16], [17]. In the northeast Atlantic context, studies indicate that cultivated seaweed environments may equal or surpass the biodiversity found in their natural counterparts by offering ‘hanging gardens’ as an alternative suspended living space for marine organisms [18], [19].

Direct observations of cultivation and header lines were conducted to assess epibiont assemblages along sections of seaweed cultivation and header lines. Our goal in these observations is to improve operational practices, focusing on balancing the need for high yields and quality with positive impacts on biodiversity. By analysing this data in the context of local physical environmental conditions, we also aim to identify the key factors driving the growth of epibionts. This is critical for creating stronger biosecurity measures, especially since the spread of non-native species on seaweed farms is a major environmental concern [20].

### 1.2 Underwater video

Seaweeds have been widely recognised for their role in creating juvenile habitats. This is mainly attributed to the protection they provide against predators and the abundance of food resources they offer, such as small crustaceans that reside in the seaweed beds [21]. However, this role is not uniform and can be influenced by various factors such as environmental conditions and human activities [22]. Most existing biodiversity studies on seaweed farms have primarily focused on benthic infauna, epifauna, and epiphytes [23]. Moreover, there is a lack of studies that explore the mobile fauna within seaweed farms, particularly in the northeast Atlantic region. This highlights the needed for more comprehensive data to understand the role of seaweed farms as juvenile habitats.

Given these considerations, our study uses underwater video to establish a baseline of the mobile fauna present beneath seaweed farms. The objective is to fill the existing knowledge gap and provide insights into the long-term ecological implications of seaweed farming, especially as commercial operations increase in scale. This will enable us to assess whether the commonly held belief that seaweeds serve as beneficial juvenile habitats holds true under varying conditions and to what extent.

### 1.3 Acoustics

In confronting an range of threats caused by humans, ranging from bycatch and pollution to habitat degradation, cetaceans such as dolphins and porpoises have become a focal point in marine conservation research [24]. Monitoring cetaceans is not only a conservation imperative but also a significant gauge for biodiversity [25]. As key indicator species, their presence or absence can reveal much about the overall health and biodiversity richness of marine habitats [26]. Moreover, with the progression of many seaweed farms in the northeast Atlantic developing from pilot to commercial scale, located increasingly in offshore environments [27], there is a critical need to better understand the potential ecological impacts on these vulnerable marine mammals [28]. In addition to these immediate concerns, our investigation aims to create a long-term record of cetacean occurrences as an indicator of biodiversity. This data will help us understand whether seaweed farming activities exert any positive or negative impacts on cetacean populations over time. Our objective is to lay the groundwork for understanding the long-term ecological interactions between seaweed farming and these protected marine mammals. By doing so, we aim to aid the development of sustainable farming practices that harmonise economic and ecological objectives. This balanced approach ensures that as we strive for commercially viable industry, we do not do so at the expense of the delicate marine ecosystems.

## 2. Materials and methods

### 2.1. Study site and farm setup

The focus of this study is on an integrated multi-trophic aquaculture (IMTA) seaweed and shellfish farm on the coast of St. Davids Peninsula, Pembrokeshire, Wales operated by Câr-Y-M r. The farm is spread across three sites, all within a 1km radius of one another. They include Porthlysgi (51.872, -5.311), Carn a Wig (51.861, -5.305) and St. Justian (51.873, -5.313) as shown in Fig. 2.A. Whilst Câr-Y-M r’s operations include the culturing of multiple shellfish species as well as seaweeds across all three sites, throughout our sampling period shellfish activities were at a small-scale and are therefore deemed to be insignificant for the purpose of our study. Porthlysgi is in a small bay south of Ramsey Sound and is open to the dominant south-westerly winds and northeast Atlantic swells. Carn a Wig and St. Justinian are inside Ramsey sound and are only open to approximately 200 km of wave fetch to the northwest. The Mean High Water (MHW) depth of all sites is between 12m and 14m. The benthic habitats can be described as sandy with large boulders at Porthlysgi and rocky reef with large boulders at Carn a Wig and St. Justinian. All sites have dominant assemblages of red seaweeds including some brown seaweeds.

The seaweed cultivation lines monitored as part of our study used a longline system with 90m - 150m cultivation lines suspended ∼2m below the surface running parallel to header lines along the surface as shown in Fig. 2.B and Fig. 2.C. Porthlysgi and Carn a Wig farms were seeded with sugar kelp (*Saccharina latissima)* in late December 2021 and harvested between May and July in 2022. Two distinct seeding methods were used at the site; Porthlysgi was seeded using the ‘twine method’ whereby 1mm thick twine with a high density of 2-10mm sugar kelp sporophytes attached, is wrapped in the cultivation line. Carn a Wig was seeded using the ‘direct-seeding method’ whereby sugar kelp gametophytes are attached directly onto 12mm thick braided rope (Langman B. V., Netherlands) using a binder solution (Hortimare B.V., Netherlands). Cultivation of seaweeds was discontinued at these sites from thereon and instead the larger St. Justinian farm was seeded with sugar kelp with the direct-seeding method in October 2022 and harvested between April and July 2023.

**Fig. 2.**
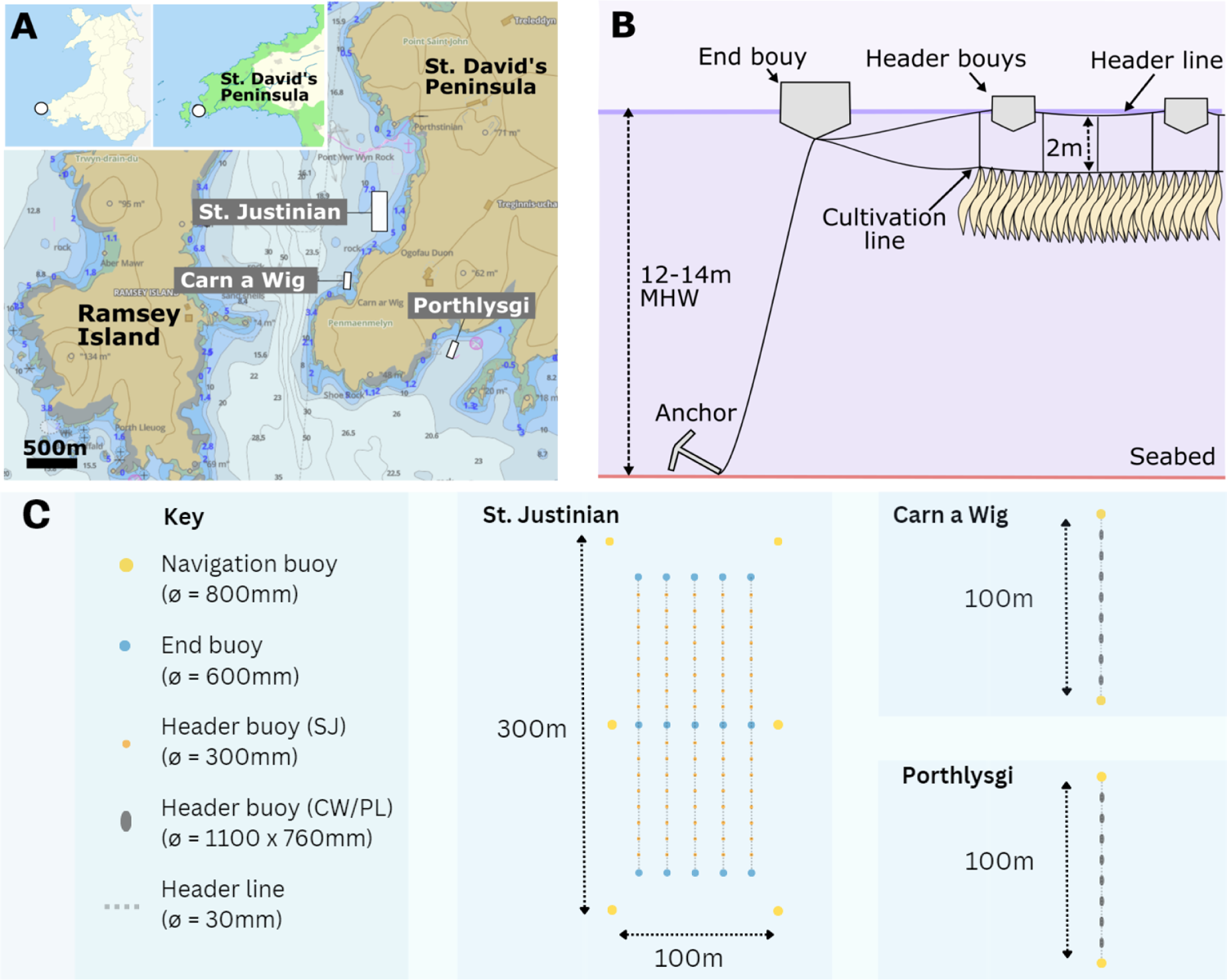
A) Map of St. David’s Peninsula highlighting the three seaweed farming areas that are the focus of this study. B) A schematic diagram of the seaweed cultivation setup showing the Mean High Water (MHW) depth in relation to the depth of the cultivated seaweeds. C) Schematic drawing of the three seaweed farming areas investigated. Drawings are not to scale.

### 2.3. Monitoring methods

#### 2.3.1 Quantifying environmental variables

At each survey site, a series of sensors were deployed to capture data on temperature, seabed irradiance, and current speed. A light sensor (Hobo pendant logger MX2202, Onset, USA) was mounted facing upwards on a buoy that was floating 0.5m up from an anchor on the seabed at ∼12m depth at each site. A control light sensor was mounted on the seabed ∼100m away from the farm at St. Justinian and Carn a Wig for measuring the relative amount of seabed shading on the seabed. An integrated temperature and current speed sensor (GrowProbe2, PEBL CIC, UK) was attached to the seaweed cultivation line at 2 m depth at each site (Fig. 3.A). Data on all three sensors was recorded every 20 minutes. Sensors were deployed beginning in January 2022 and were retrieved and swapped out with a replica of the sensors each month up to 1st July 2023. Water samples were collected monthly at each site by filling a 500ml sample bottle at 0.5m depth from the side of the monitoring vessel. Water samples were analysed within 3hrs of sample collection for pH and Salinity (PCE-PHD1, PCE Instruments, Germany).

#### 2.3.2. Direct observations of cultivation lines

Epibiont assemblages growing on the rope of the header line and on the cultivation line, including the cultured sugar kelp canopy were quantified along 0.5m sections by pulling the lines onto the research vessel and cutting all seaweeds off at ∼2cm from the line. At each of the sites, two locations were sampled on opposite ends of the long-line, approximately 90m apart. Samples were stored in a closed container and brought back to land for analysis. For each 0.5m sample batch 10 individuals were selected at random for which the area coverage and taxa of epibionts directly on the line and on a single side of the kelp were estimated visually.

#### 2.3.3. Baited Remote Underwater Video (BRUV)

Baited Remote Underwater Video systems, or BRUVs, are effective instruments for studying changes in the distribution, behaviour, and relative abundance of underwater fauna. The commonly used metric, Nmax, derived from BRUVs, quantifies the relative abundance of fish observed in underwater videos by counting the peak number of fish seen at any single moment, eliminating the risk of counting the same fish twice [29]. Despite the turbid waters around the UK, BRUVs are emerging as a useful tool to assess the biodiversity of coastal habitats [29], [30], [31], [32]. In the present study the BRUV setup includes a large-battery power video camera (SubCam2, PEBL CIC, UK) fixed onto a galvanised steel frame (Fig. 3.A). Attached to this frame is a pole, extending approximately 90cm from the camera, with a bait bag filled with brown crab meat (∼500g, defrosted 2-3hrs before deployment), allowing for effective attraction and observation of marine life.

All BRUV sampling was conducted during the daytime using either a small research vessel or a work barge. The deployment duration used in this investigation was 1h, as this was deemed suitable for assessing fish communities using BRUV [29]. All deployments were in a depth range of between 8 and 14m and were placed away from cliff edges to avoid the presence of any edge effects. To ensure consistent data collection, the BRUV was isolated from surface water movement by using a combination of floating and leaded rope between the surface and anchor. Due to the difficulty in working in coastal waters of high turbidity, footage was classified in terms of quality. Only footage where the observer could clearly see the bait bag (90 cm distance) and where organisms could be identified at least to the family level was analysed.

#### 2.3.4. Passive Acoustic Monitoring (PAM)

Acoustic monitoring is a useful tool for examining fine scale changes in distribution, behaviour, and relative abundance of vocalising cetaceans [33]. Static omni-directional Porpoise Detectors, known as PODs (Chelonia Ltd, UK) have been widely used to study the relative abundance, behaviour, habitat use, and distribution of cetacean species in the west Wales region [34], [35], [36], [37]. PODs detect echolocation click trains, and both dolphins and porpoises produce these high frequency clicks for navigation as well as prey detection and discrimination. Acoustic detection rates are thought to relate to the rate of occurrence of an echolocating species and may give an indication of relative abundance [38]. PODs are self-operating devices that detect dolphin and whale echolocation sounds. They have a hydrophone linked to an amplifier and filters set to specific frequencies. These devices record specific high-pitched sounds and can work continuously for 4-6 months, influenced by sea temperature and settings. A comprehensive description of the POD units used in this study (F-POD) can be accessed at www.chelonia.co.uk/fpod_home_page. Similarly to the BRUV setup, the POD was isolated from surface water movement by using a combination of floating and leaded rope between the surface and anchor, as well as an intermediate anchor.

Acoustic data was extracted using F-POD.exe software (version 0.91; Chelonia Ltd, UK), which employs an integrated train detection algorithm to filter raw click data, identify cetacean click trains, and estimate the probability of their occurrence by chance from non-train producing sources (e.g., rain or boat propellers). The software subsequently categorises the click trains based on species and the likelihood of cetacean origin. To minimise the inclusion of false positive detections, only data classified by the algorithm as highly likely to be of cetacean origin were utilised. Though this approach resulted in some missed detections, given the substantial number of detections for both species at the study site, no additional false positive analysis was deemed necessary. The presence of dolphins and porpoises at each site was described by exporting the number of detection positive minutes (DPM) per hour and per day for both species.

### 2.4 Sampling schedule and locations

The schedule of all monitoring activities, including their specific time frames and any gaps in data, can be found in Fig. 3.A, while Fig. 3.B offers detailed information on the exact locations, accurate to within 10 meters, where each monitoring method was deployed. Visual observations were carried out monthly at Porthlysgi and Carn a Wig from April 2022 to June 2023. At the St. Justinian site, construction and modifications to the farm site meant visual observation were only possible from August 2022 to June 2023. BRUV units were deployed monthly from April 2022 to July 2023, at Porthlysgi ad Carn a Wig. Deployments were carried out at St. Justinian site in September 2022, April, and June 2023 for additional comparison. Data gaps correspond to periods where units could not be deployed due to ongoing marine works. The PAM unit was deployed for four periods spanning a total of 21 months near to St. Justinian at a depth of 9m below mean high water. Data was successfully captured for 162 days across 3 deployment periods in summer/autumn 15/08/2022 – 21/09/2022 (38 days), winter 11/11/2022 – 16/12/2022 (36 days), and spring 03/04/2023 – 22/05/2023 (50 days). Data gaps correspond to periods where the probe was removed for maintenance or had been removed due to nearby marine works. We placed environmental sensors and collected water samples every month at Porthlysgi from April 2022 to July 2023. During the same time frame, we also carried out monthly measurements at Carn a Wig and St. Justinian. However, we stopped these measurements at Carn a Wig in October 2022 as it became evident that water quality measurements were identical for the two sites.

**Fig. 3.**
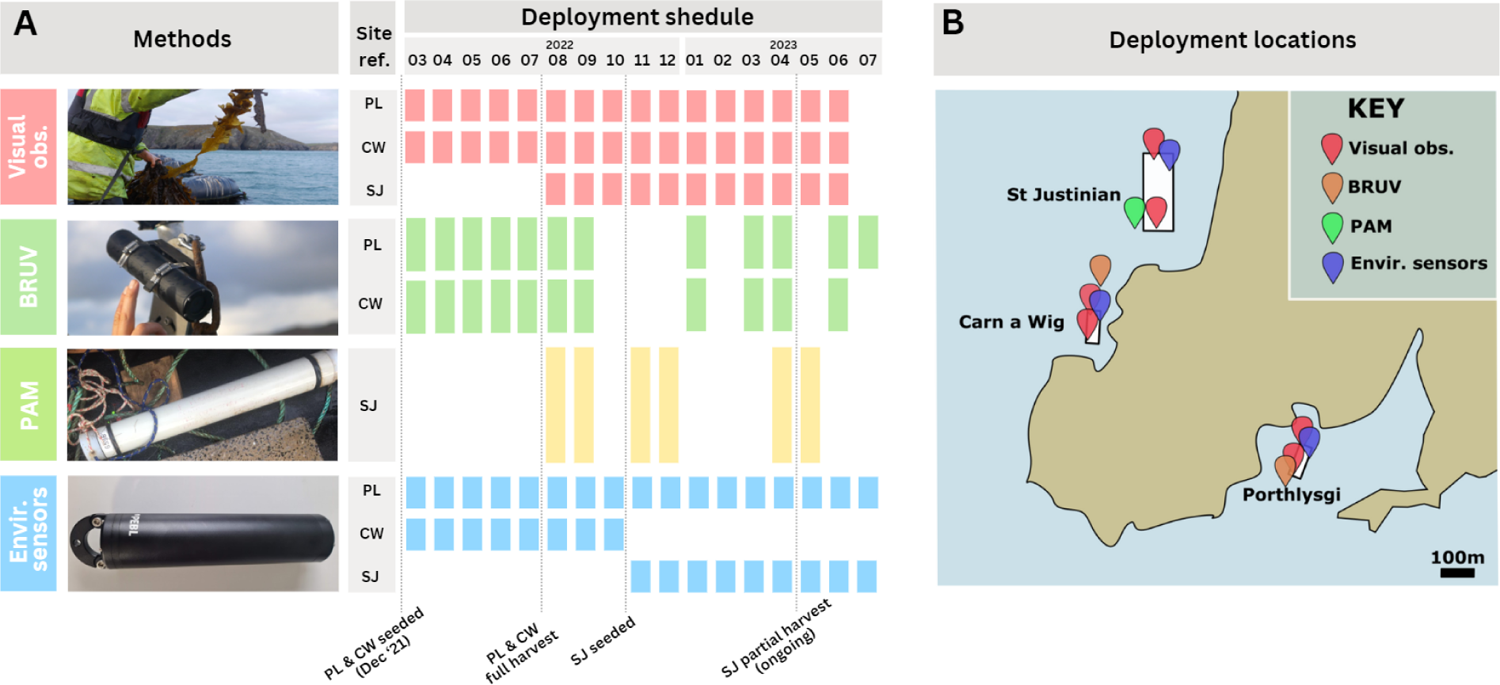
A) A timetable showing when each monitoring task was performed is provided. Dotted vertical lines on the chart indicate important actions that took place at the seaweed farms. The locations of these farms are abbreviated as follows: PL stands for Porthlysgi, CW for Carn a Wig, and SJ for St. Justinian. B) A map showing the location of where monitoring activities were carried out on each of the seaweed farming sites.

## 3. Results

### 3.1 Quantifying environmental variables

Throughout the monitoring period, environmental parameters at Carn a Wig and St Justinian were almost identical. Environmental data for these two sites is therefore presented under a single site heading (i.e. St. Justinian / Carn a Wig). Detailed variations in physical environmental conditions (temperature, irradiance and current speed) over the course of the study are shown in Fig. 4. The measurements taken in January and July served as inflection points for all parameters, signifying a plateau and subsequent reversal of physical environmental trends. Data from these two months were therefore considered to give the best summary of the minimum and maximum environmental conditions at the sites, as detailed in Table 1.

Water quality related environmental conditions, however, did not show clear annual trends. PO₃⁻ levels remained consistent across sites and months, hovering around 0.06 µM. Similarly, NO₃⁻ + NO₂⁻ concentrations stayed stable, with only slight July elevations. All sites maintained a similar pH, though it marginally rose to around 8.39 in July. Likewise, salinity in January was consistent at about 29.6 psu, but July saw a slight increase, especially for St. Justinian/Carn a Wig at 31.1 psu.

**Table 1.** Summary of environmental conditions at each study site. “Mean SST” is the mean sea surface temperature measured at 2m depth across each month. “Mean irradiance” is the mean seabed irradiance measured at a depth of 12m for Carn a Wig and Porthlysgi for the period of each month. “Mean current speed” is the mean current speed measured at a depth of 2m for the period of each month.

| Site | Month in 2023 | Mean SST (°C) | Mean seabed irradiance (lux) | Mean current speed (m/s) | $\text{PO}_4^{3-}$ ($\mu\text{M}$) | $\text{NO}_3^- + \text{NO}_2^-$ ($\mu\text{M}$) | pH | Salinity (psu) |
| --- | --- | --- | --- | --- | --- | --- | --- | --- |
| St. Justinian / Carn a Wig | January | $9.64 \pm 0.9$ | $20 \pm 43$ | $0.33 \pm 0.36$ | $0.04 \pm 0.01$ | $0.23 \pm 0.02$ | $8.32 \pm 0.02$ | $29.6 \pm 0.5$ |
| Porthlysgi | January | $9.68 \pm 0.9$ | $22.9 \pm 55$ | $0.41 \pm 0.39$ | $0.04 \pm 0.01$ | $0.23 \pm 0.02$ | $8.32 \pm 0.02$ | $29.6 \pm 0.5$ |
| St. Justinian / Carn a Wig | July | $15.33 \pm 1.5$ | $121 \pm 269$ | $0.25 \pm 0.29$ | $0.06 \pm 0.01$ | $0.27 \pm 0.03$ | $8.39 \pm 0.03$ | $31.1 \pm 0.2$ |
| Porthlysgi | July | $18.55 \pm 4.6$ | $434 \pm 950$ | $0.14 \pm 0.12$ | $0.06 \pm 0.01$ | $0.27 \pm 0.04$ | $8.39 \pm 0.05$ | $31.1 \pm 0.3$ |

January saw similar temperature patterns at both sites, with minimal day-night variation and an average temperature around 9.8°C. As summer approached, Porthlysgi recorded higher temperatures, ranging from 14°C to 28°C, compared to Carn a Wig’s steadier range of 15.5°C to 16.5°C. Porthlysgi also experienced more pronounced daily temperature fluctuations, reaching up to 8.5°C, in contrast to Carn a Wig’s less than 1°C variation.

Initial irradiance measurements in January were comparable at both sites. However, with the onset of summer, Porthlysgi showed higher irradiance levels, with July readings 4-10 times greater than at Carn a Wig. This indicated a marked difference in water column transparency between the sites during this period.

In January, Porthlysgi recorded slightly higher current speeds than Carn a Wig. By July, Carn a Wig’s speeds remained consistent, while Porthlysgi’s dropped significantly. Carn a Wig displayed a consistent semi-diurnal cycle in current speeds, minimally affected by storms. Porthlysgi’s speeds, however, showed partial tidal synchronisation and were more influenced by storm events, with a notable increase during a storm in early January. In July, Porthlysgi’s current speeds were very low, occasionally near the detection limit, contrasting with Carn a Wig’s clear semi-diurnal patterns.

**Fig. 4.**
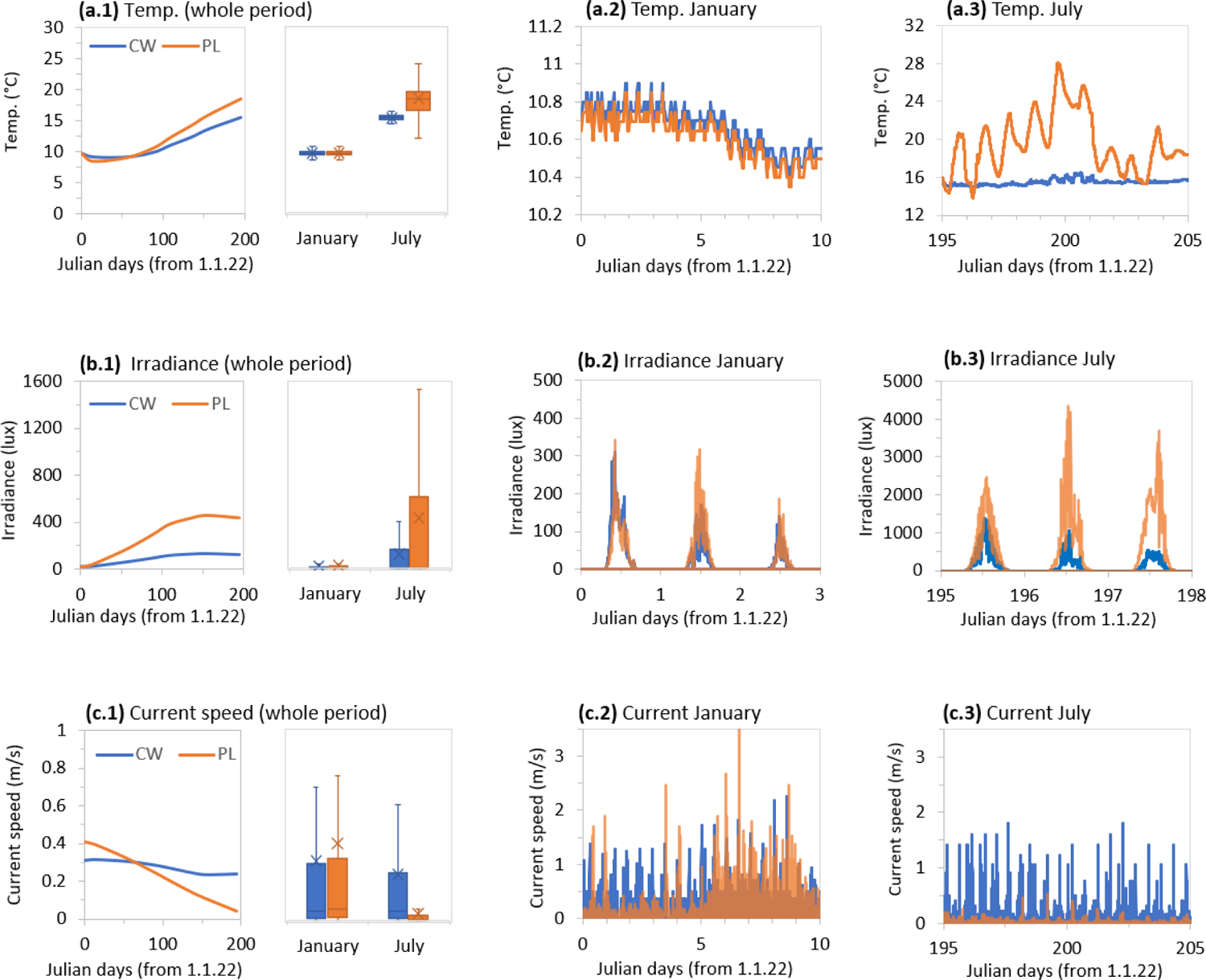
Sea surface temperature, seabed irradiance and current speed recorded during the study period. Blue plot is data for Carn a Wig (CW) and orange plot is for data at Porthlysgi (PL). (a.1 – c.1) Line plot and box-whisker plot of sea surface temperatures, seabed irradiance and current speed recorded over study period and for 31 days in January and 31 days in July 2022 respectively. Outlier points have been omitted from box-whisker plot (a.2 – c.2). Line plot of sea surface temperatures, seabed irradiance and current speed recorded for 10 days, 3 days and 10 days from 1.1.22 respectively (a.3 – c.3). Line plot of sea surface temperatures, seabed irradiance and current speed recorded for 10 days, 3 days and 10 days from 1.1.22 respectively.

### 3.2 Direct observations of header and cultivation lines

#### 3.2.1 Composition of seaweed species on header lines

The composition of seaweed species settled on the header lines of the seaweed farm varied across the sites Porthlysgi, Carn a Wig, and St. Justinian, reflecting the variable environmental conditions and life cycles of seaweeds at each location. Fig 5.a shows the header line coverage at Porthlysgi throughout the monitoring period. Oarweed (*Laminaria digitata*) and a diverse range of red algae (including *Chondrus crispus, Osmundea pinnatifida, Dilsea carnosa*) dominated the line coverage. A notable increase in Oarweed was recorded during autumn months, displacing many of the red algae. Carn a Wig’s header line showed a more balanced assemblage, with red algae prevalent throughout most of the year (Fig 5.b). Conversely to Porthlysgi, Oarweed did not proliferate during the winter period, but instead Sea lettuce (*Ulva sp.*) became more prominent. Both Porthlysgi and Carn a Wig showed a clear peak in Furbellows (*Sacchariza Polyschides*) during spring and autumn months. Although St. Justinian was only monitored for 4 months, the naturally settled assemblage was largely dominated by red algae, with consistent coverage throughout the monitoring period.

**Fig. 5.**
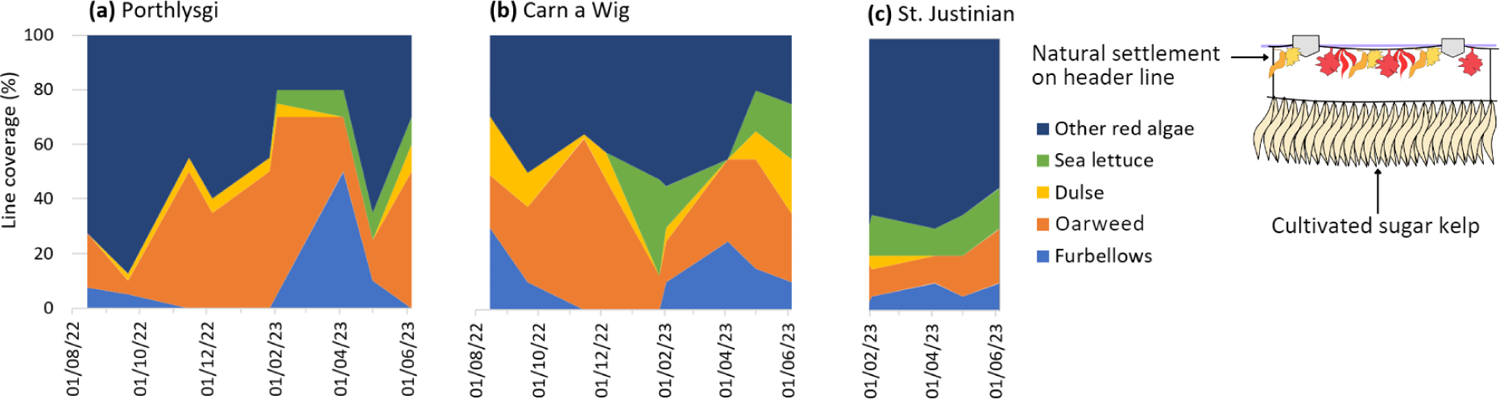
Seasonal line coverage percentages by seaweed species on the header lines at (a) Porthlysgi, (b) Carn a Wig, and (c) St. Justinian from August 2022 to March 2023. The coverage is represented by various colours, each corresponding to a different species, with the inclusion of Other red algae such as Chondrus crispus, Osmundea pinnatifida and Dilsea carnosa commonly found at these depths.

#### 3.2.1 Morphology of cultivated sugar kelp and corresponding seabed shading

The morphology and growth dynamics of cultivated kelp were highly variable across all study sites, particularly during the summer months. The weight per metre of cultivation line (yield) recorded at both sites showed a similar trend, reaching maximum lengths on day 135 (Fig. 6.a). However, in the following 20-day interval, a sharp increase in weight was recorded at Porthlysgi, from 5.2 kg/m to 13.5 kg/m, with a large standard deviation (±7.6 kg/m). Meanwhile, at Carn a Wig, the length showed a linear increase with a relatively consistent standard deviation during the same period. At Porthlysgi, sugar kelp lengths grew faster than at Carn a Wig, reaching a length of 140 cm approximately one month earlier (Fig. 6.b). The peak length recorded at Porthlysgi (169 ± 4 cm) was also significantly larger than at Carn a Wig (141 ± 5 cm), sampled on day 135. Subsequent measurements showed a slight decrease in length at both sites, with the standard deviation in measurements steadily increasing beyond the peak length measurement. The maximum width of kelp blades recorded at both sites followed a similar trend in line with length measurements up to the peak length sampling (Fig. 6.c). However, in the following 20-day interval, a sharp step increase in width was recorded at both sites, with Porthlysgi increasing from 4.5 ± 0.4 cm to 10.4 ± 1.5 cm and Carn a Wig increasing from 4.2 ± 0.3 cm to 9.3 ± 1.2 cm. Beyond this step increase, both sites continued to increase but at a steadier rate.

The net growth rate was determined for both sites by taking the difference between subsequent samplings and dividing by the number of days between them (Fig. 6.d). The peak growth of kelps was observed in the mid-spring to early summer period, between day 90 and 135, with net growth rates peaking at 2.1 ± 0.2 cm/day on day 108 for Carn a Wig and 2.6 ± 0.3 cm/day on day 89 for Porthlysgi. Both sites showed a subsequent decrease in net growth down to a negative value, with Porthlysgi showing a sharper and more negative growth than Carn a Wig. In the final sampling, net growth was net positive once more, but only marginally, at 0.5 ± 0.1 cm/day at both sites.

During the initial stages of development, kelp blades exhibit a tapered tip that gradually becomes fragmented as the blade reaches maturity. Reproductive cells are primarily concentrated near the stipe of the plant, resulting in a conveyor-belt-like growth pattern in which new cells emerge at one end while old cells disintegrate at the other. By analysing the relationship between the maximum blade width and the total plant length, it is possible to estimate the amount of kelp material lost through shedding when specific blade tip widths are observed (Fig. 6.d) making the kelp blade tip wider and wider.

**Fig. 6.**
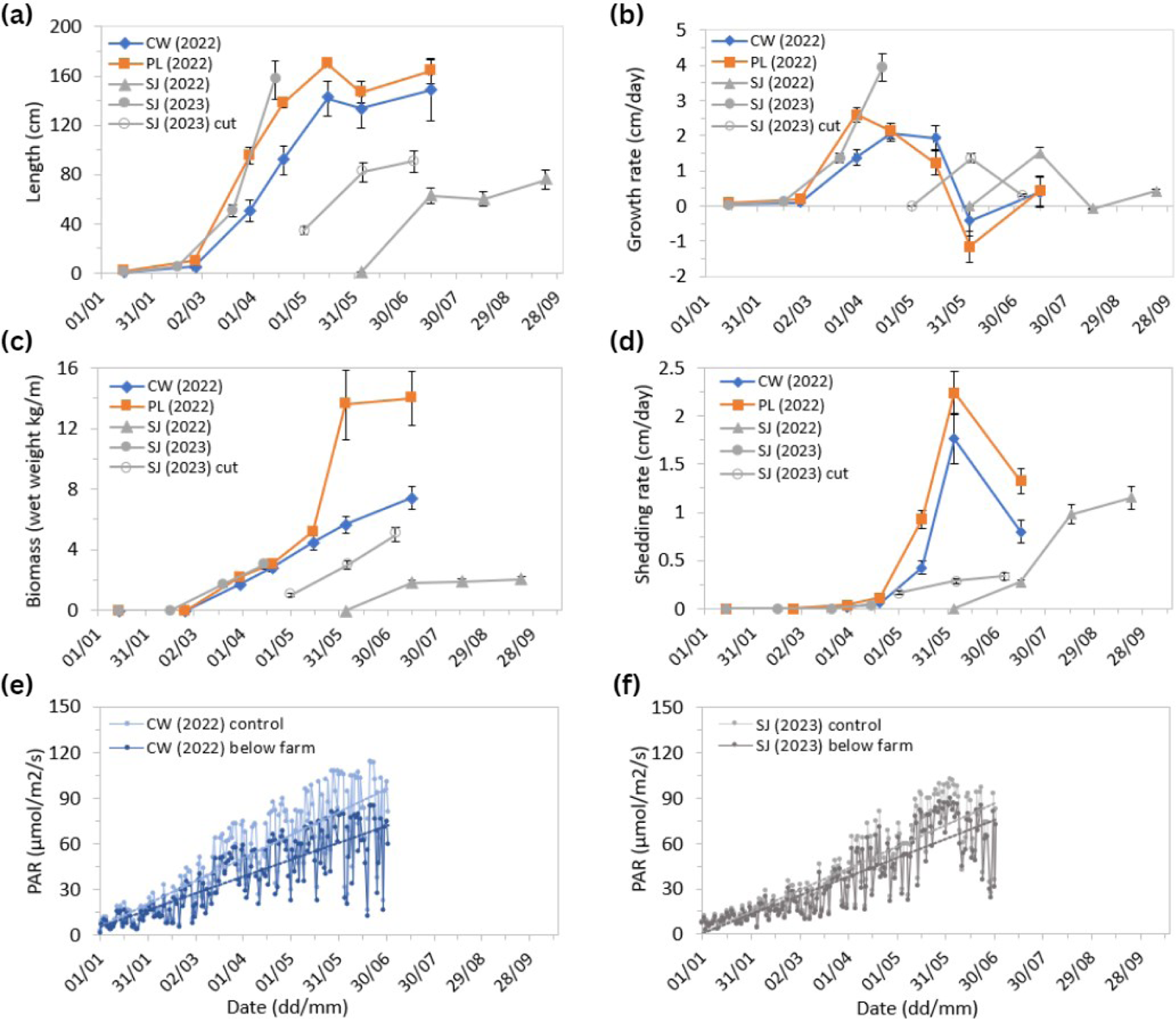
Kelp morphology and growth variables recorded throughout 8 sampling periods from seeding to harvest. Blue plots are datapoints for Carn a Wig (CW) and orange plot is for data at Porthlysgi (PL). Each data point represents an average of multiple measurements as described with standard deviation as error. (a) Kelp length (from holdfast to blade tip) recorded from 20 individuals at each site (10 at each end of site). (b) Net growth rate calculated from length measurements in (a). (c) Biomass or kelp wet weight per meter (when cut along holdfast 2-5cm from cultivation rope) recorded from 2 samples at each site (1 sampling at each end of site). (d) Shedding rate (tip fragmentation) approximated from blade-tip width measurements from 20 individuals at each site (10 at each end of site). (e) Photosynthetically Active Radiation (PAR) recorded below the seaweed farm during daylight hours (averaged for each day) compared to a control site (away from seaweed farm) in 2022 and (f) in 2023.

Therefore, by measuring kelp blade tip width an indirect measure of the shedding rate is obtained. Plotting the shedding rate reveals that shedding initiation occurs in late spring to early summer on day 89 at both locations. Subsequently, the shedding rate undergoes a rapid acceleration, reaching a peak of 3.6 ± 0.9 cm for Carn a Wig and 2.8 ± 0.8 cm for Porthlysgi, both on day 155. During the final sampling, the shedding rate was notably lower for both sites, and a distinct difference between the two locations emerged, with Carn a Wig displaying a rate of 2.3 ± 0.8 cm and Porthlysgi demonstrating a substantially lower rate of 0.6 ± 0.5 cm. The total productivity of kelp can be assessed by combining its net growth rate and shedding rate. This metric serves to examine the relationship between environmental factors and the kelp’s growth and morphology and is further discussed in subsequent sections.

Light measurements at 12m depth showed significant variation in light penetration at the seabed beneath the seaweed farm compared to a control site located away from the farm (Fig. 6.e and Fig. 6.f). In 2022, the PAR levels beneath the farm at Carn a Wig were consistently lower than the control, suggesting that the kelp canopy and associated buoys effectively reduced light availability to the benthos. This shading effect was less at St. Justinian in 2023 with approximately 60% less shading than observed at Carn a Wig in 2022.

#### 3.2.2 Biofouling of cultivated sugar kelp

Four types of biofouling organisms (sea fur, bleaching, filamentous algae, and bryozoans as shown in Fig. 7.a-d) were found to be dominant amongst the cultivated kelp canopies. These were monitored monthly at both sites but were only observed in significant quantities during the final 3 samplings before harvest. Overall, Porthlysgi had a greater percentage blade area covered by fouling organisms than Carn a Wig, with up to 73.5% biofouling blade coverage by the final sampling (Fig. 7.e). While biofouling did occur at Carn a Wig, the mean blade cover at the final sample was 23%, with filamentous algae making up the highest cover up to 11% in the final sampling.

Filamentous algae, primarily *Ectocarpus spp*. were located on the distal tips of the kelp covering 5.5% at both sites on day 135 and continued to propagate on the tips of blades through to the final sampling on day 195 with 11% and 26.5% blade coverage at Carn a Wig and Porthlysgi respectively. The colonial organisms, sea fur (*Obelia geniculata*) and bryozoans (*Membranipora membranacea*), also spread rapidly to high levels of cover by the final sampling with 17% and 13% coverage at Porthlysgi and 2% and 7% coverage at Carn a Wig respectively. Bleaching of kelp blades was mostly observed only in the final sampling with 4% at Carn a Wig and 17% at Porthlysgi.

**Fig. 7.**
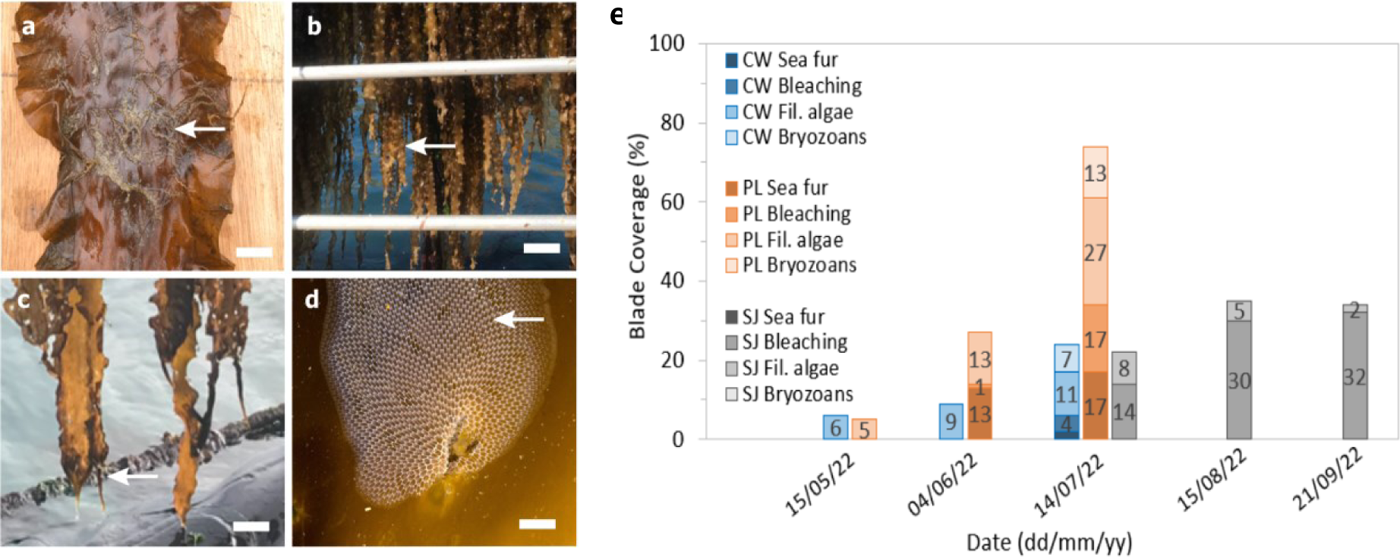
(a-d) Photographs of epiphytes and fouling on cultivated kelps taken throughout the final 3 sampling periods before harvest. White arrows point at fouling features. (a) Sea Fur (Obelia geniculata) on the surface of a kelp blade. Scale bar: 2cm. (b) Bleaching (discoloration) on the tips of kelp blades. Scale bar: 20cm. (c) Filamentous algae (Ectocarpus spp.) on distal tips of kelp blade. Scale bar: 10cm. (d) Bryozoans (Membranipora membranacea) on surface of kelp blade. Scale bar: 0.5cm. (e): Biofouling on cultivated kelps recorded throughout the final 3 sampling periods before harvest. Blue, orange and grey columns show data for Carn a Wig (CW), Porthlysgi (PL) and St. Justinian (SJ) respectively. Each column shows culminated mean blade coverage of the biofouling organisms shown in the photographs in (a-d).

### 3.3. Baited Remote Underwater Video (BRUV)

Our study identified 13 taxonomic groups, 10 of these taxa were observed at Porthlysgi (sand and occasional boulders) and 9 at Carn a Wig (reef and boulders). Of these 13 taxa, 13 different families and 14 different species were identified. The average relative abundance of mobile fauna (Total Nmax or sum of all individual species’ Nmax in each 1h sample) was 9.9 ± 5.7 SD. Whiting-pout (*T. luscus*) were the most abundant species at both sites with an average Nmax of 2.8 ± 2.3 SD per sample at Porthlysgi, and average Nmax of 3.9 ± 4.1 SD per sample at Carn a Wig. The second most abundant species were greater spotted dogfish (*S. stellaris*) at 2.0 ± 2.3 SD at Porthlysgi and pollack (*P. pollachius*) at 2.4 ± 2.9 SD at Carn a Wig. Whiting-pout had the highest individual species Nmax with 13 individuals in a sample, followed by pollack at 9 individuals in a sample. Both these values were achieved near the end of the 1h deployment.

**Table 2.**
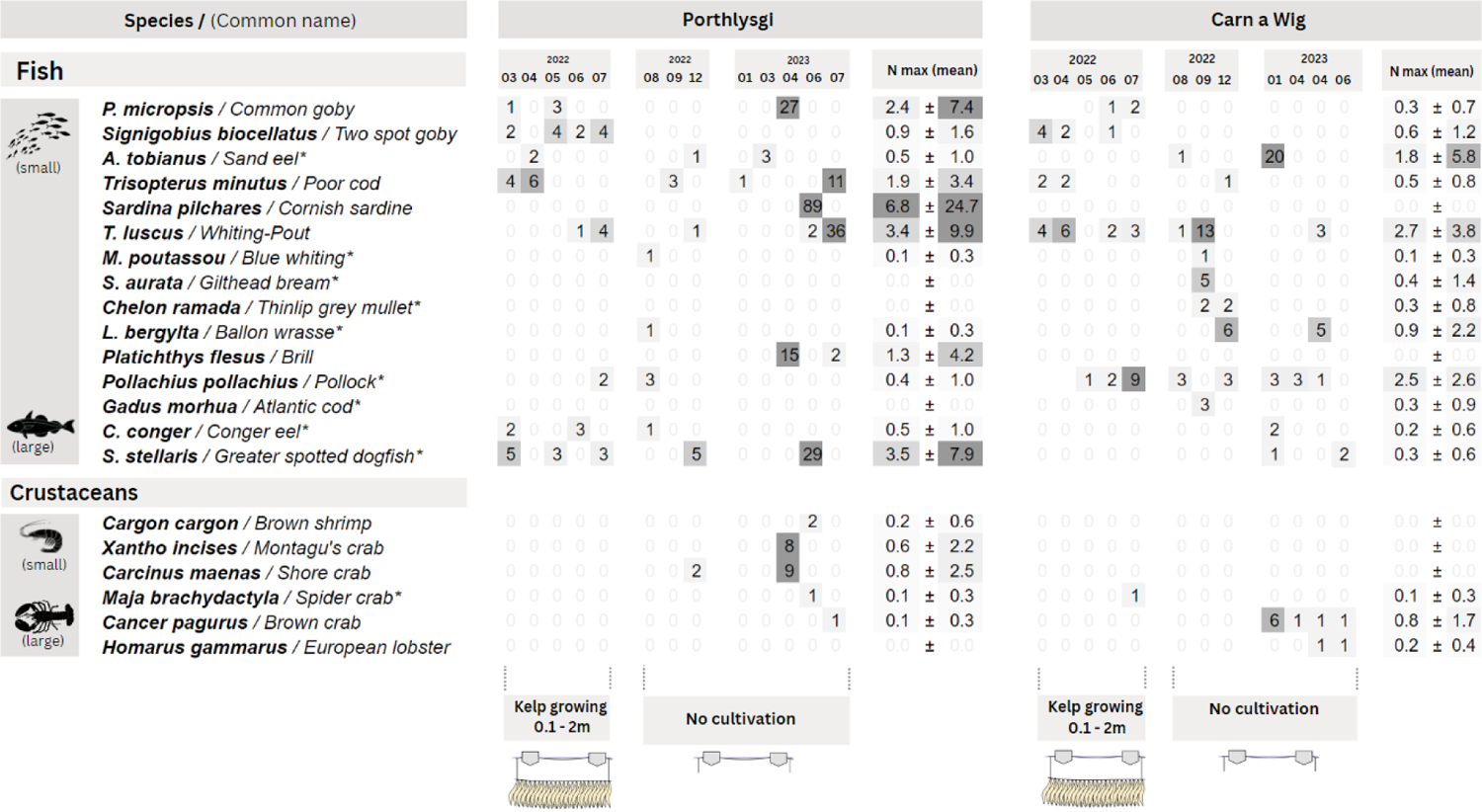
Mean (±SD) relative abundance (Nmax) and length for each species within Porthlysgi (PL) and Carn a Wig (CW) habitats using BRUV. Species denoted with a * are commercially important species in the UK.

### 3.4. Passive Acoustic Monitoring (PAM)

Figure 8.a shows the detection patterns for dolphins and porpoises during the study period. Throughout, both species showed variability in their detections, with neither species being exclusively detected during any period. A notable surge in detections for both species occurred from 28th November to 16th December 2022. In the final monitoring period, from 8th to 22nd May 2023, harbour porpoise detections significantly outweighed those of dolphins. While fluctuations are present, a consistent seasonal trend is not obvious. Nonetheless, the summer/autumn and winter periods indicate a close alignment in the detection positive minutes per day (DPM/day) for both species.

When evaluating the mean DPM per hour daily within each monitoring period, porpoise detections consistently exceed those of dolphins throughout the diurnal cycle. In the summer/autumn period, peak detections for both species predominantly occurred during the night, from 21:00–06:00. Detections sharply declined after sunrise and remained minimal during daylight, before surging around 19:00, coinciding with sunset. The winter months mirrored this diurnal pattern, albeit less pronounced. By spring, a clearer diurnal pattern emerged again. While porpoise detections were not entirely absent during spring, as previously observed in summer, they were significantly reduced during daylight hours.

An integral part of our study was measuring the residency time of dolphins and porpoises near the seaweed farm. For harbour porpoises, the residency time ranged between 8 to 14 minutes. In contrast, dolphins exhibited slightly longer residency times, varying from 11 to 16 minutes. These relatively short durations suggest that both dolphins and porpoises were transient visitors to the seaweed farm area, likely passing by without engaging in significant feeding activities at the site. The residency times also did not display significant variations or discernible trends over time. This pattern remained consistent across different months and even on a week-to-week basis.

**Fig. 8.**
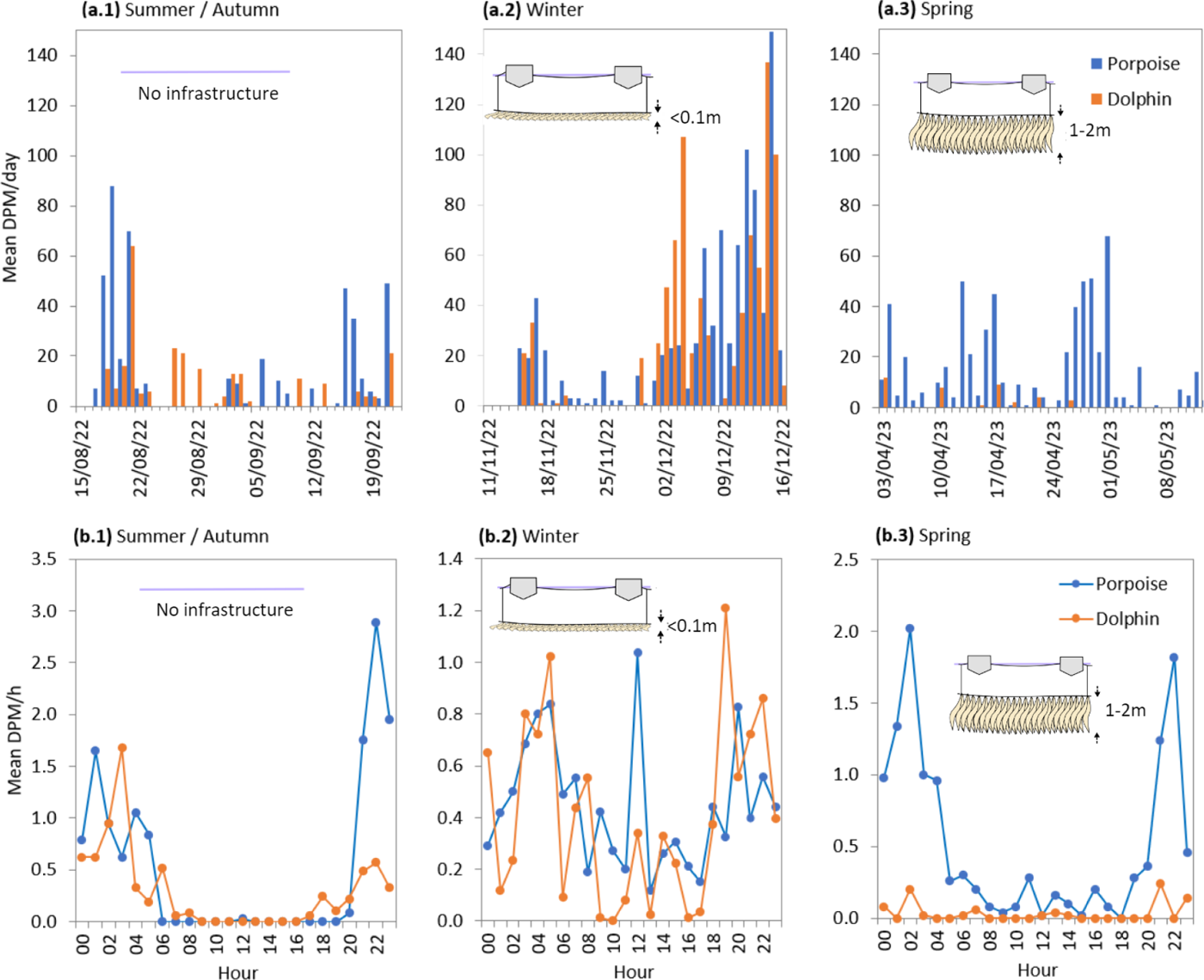
(a.1-3) Daily variation in porpoise and dolphin mean DPM per day over the three monitoring periods. The insets show the configuration of the seaweed farm at St Justinian, with no cultivation line in summer/autumn, and the seaweed canopy at a length of <0.1m and 1-2m in winter and spring, respectively. (b.1-3) Diurnal variation in DPM per hour for each of the same three monitoring periods.

## 4. Discussion

### 4.1. Direct Observations

#### 4.1.1. Interaction between environmental factors

Our study noted distinct environmental patterns due to the specific geographical and climatic influences of each site. Porthlysgi, situated in a bay with relatively low tidal currents, faces direct impact of storm events due to its exposure to prevailing weather fronts from the southwest. These conditions lead to a drop in sea surface temperatures and a decline in water clarity, as the storms stir up sediment and mix the surface water with deeper, colder waters. The changes in sea surface temperature and turbidity at Porthlysgi are more dramatic, reflecting the direct impact of these storm events. In contrast, Carn a Wig has a more protected position, shielded from prevailing storms, and influenced by strong tidal currents, which maintains more consistent and stable environmental conditions.

These environmental differences have implications for seaweed farming. At Porthlysgi, the variable conditions may require for substantial moorings and large line spacing that can withstand the repeated storm impacts and prevent lines from becoming entangled with one another. In contrast, Carn a Wig’s cooler, stable environment could allow for seaweed farming throughout the summer, avoiding fouling issues prevalent at Porthlysgi from mid-year. Although Carn a Wig endures relatively high tidal current speed, the likelihood of storm damage is significantly reduced as North-westerly storms are much less frequent and bring with them smaller wave heights.

Regarding the broader ecological implications, Porthlysgi’s dynamic conditions, with their higher storm swells, might hinder the establishment of stationary species due to the physical disturbance of the seabed, as observed in other regional studies [39]. In contrast, Carn a Wig, with its lower exposure to extreme weather events, provides a more conducive environment for such species to establish.

Overall, our findings underscore the importance of site-specific approaches in seaweed farming. In dynamic sites like Porthlysgi, strategies should focus on cultivating resilient species or adjusting harvest times. Conversely, at stable sites like Carn a Wig, leveraging consistent conditions can extend the growing season and enhance yield.

#### 4.1.2 Composition of seaweed species on header lines

The natural settlement observed on header lines at Porthlysgi fluctuates significantly from winter to summer, attributed to the higher storm exposure and significantly warmer summer waters. Carn a Wig, however, maintains a more uniform species presence year-round, due to its sheltered location, shielding it from such environmental extremes. Observations at St. Justinian show that the seaweed composition is still in its early stages of development. Being a newly established farm, the expected diversity has not yet fully materialised. Over time, as the farm matures, a more diverse assemblage is anticipated, reflecting the patterns observed at more established sites [40].

#### 4.1.3 Influence of light on growth and morphology

At each site, all morphological parameters showed an increase during the culture period, in line with an increasing light availability. The growth at Porthlysgi was consistently ahead of that at Carn a Wig by approximately 30 days, even though both sites were seeded with identical seed stocks within a two-day interval. This discrepancy can be attributed to the higher sedimentation at Carn a Wig, which resulted in diminished light availability due to an increased turbidity and sediment deposition on the kelp seedlings’ surfaces. The latter was backed by observations of a thin film of sediment on juvenile kelp blades during late January and February at Carn a Wig. Similar studies in the North Atlantic have found that factors such as increased sedimentation and reduced light availability, can stress kelp species and alter their growth patterns [41], [42].

It is important to highlight that the day light duration varies between the two sites, especially during the early hours of the day. When the sun is close to the horizon as Carn a Wig is obstructed by an approximately 20m-tall cliff roughly 40m to the east, whereas Porthlysgi, with a cliff face to the north, has direct sunlight throughout the day. This effect is also likely to add to the early (winter) season growth discrepancy between the two sites where the sun remains low on the horizon.

#### 4.1.4 Influence of temperature on growth and morphology

On exposure to temperatures above their optimal range, sugar kelp can undergo thermal stress, which can stimulate adverse effects such as diminished growth rates, compromised photosynthetic efficiency, and cellular damage [43]. Variations in sea surface temperature, particularly during warmer months, have been shown to modulate stress and energy budgets, potentially accounting for a portion of the discrepancies in growth [44]. This is supported by our study when examining temperatures at peak productivity at Carn a Wig (peak at March 30^th^) and Porthlysgi (peak at June 4^th^), which are both within the range of 12-13°C. Similar observations have previously been made in the UK, where sugar kelp showed optimal growth at temperatures close to 12°C, beyond which productivity began to slow down [45].

This highlights that the rising sea temperatures due to climate change present a critical challenge for seaweed. To navigate these challenges, strategies such as selective breeding for more heat-resistant kelp varieties [46], timing adjustments in cultivation to align with optimal growth temperatures, and exploring deeper waters with more stable temperatures will be crucial [47].

#### 4.1.5 Influence of current speeds on growth and morphology

Prior research has shown minimal impacts of changes in water motion on growth rates [48], suggesting that current speeds are more likely to indirectly influence growth by affecting turbidity and sea temperature. Previous investigations examining kelp farm yields in relation to current speed have centred on various exposure types (e.g. sheltered, current-exposure or wave-exposure), leading to unclear results and making it challenging to derive general conclusions. For instance, highly wave-exposed environments were associated with lower yields [49], while in another study higher yields were reported at both wave-exposed and sheltered sites compared to current-exposed locations [52]. Typically, seaweeds exhibit narrower blades in current-exposed habitats and wider blades in sheltered environments [50], [51]. The presence of narrower blades at the current-exposed site of Carn a Wig in the present study supports this pattern.

The effects of current-exposure on the shedding of kelp blade tips have previously been shown to be significant [52]. High current speeds can cause the blade tips to break off, resulting in reduced photosynthetic capacity as well as a decrease in the quality of the blade itself due to physical damage. These effects are supported by our study as a significantly higher shedding rate was observed in the higher current-exposed site, as shown by Fig. 6.d.

Increased frequency, severity, and off-season occurrences of storms due to climate change will significantly impact the physical dynamics within kelp farming areas. Impacts are likely to lead to increased blade tip shedding and in extreme storm events, complete dislodgement at the holdfast. This necessitates for adaptive strategies in farm management similar to those proposed for increased water temperatures, including the installation of adjustable depth cultivation lines to modulate the depth of seaweed grow-lines away from the high-energy sea surface, coupled with selective breeding programs that focus on developing kelp strains with enhanced resilience to mechanical stress from water motion [47].

#### 4.1.6 Influence of water quality and grazers on growth and morphology

Our study found that water quality variables such as nutrient availability, pH, and salinity were almost identical at both sites and did not show significant variation throughout the year. As a result, they were assumed to be insignificant in accounting for any differences between the sites. Furthermore, all water quality parameters were like the seasonal average values for the Celtic Sea, indicating limited influence from the coastal environment.

Previous studies have shown that biotic factors such as grazing by herbivores, competition with other organisms for resources, and the presence of symbionts, parasites, or pathogens can significantly affect the growth dynamics of sugar kelp [53]. While significant biofouling was observed at both sites during the June and July samplings, there was no significant correlation with the morphology or growth of the plants during this period, suggesting that the influence of biotic factors on growth was minimal. Instead, it is more plausible that the elevated temperatures during these summer months played a dominant role in reducing kelp growth, overriding the potential effects of biotic stressors.

#### 4.1.7 Human factors effecting kelp growth and morphology

The differences in growth trends observed at the two farm sites could potentially be attributed to human error during the process of deploying the seedlings. Although the same methodology was applied at both sites, there were several variables in the seeding process that could not be controlled, such as the tension and pitch of the seeded twine coil, as well as the density of kelp seedlings on the twine. These factors may have significant effects on the density of the kelp canopy and, subsequently, on its morphology as reported in previously at seaweed farms in the UK [54]. Variations in water depth and light availability, caused by slack in the farm’s design, might have also affected growth patterns. However, random sampling and a large sample size likely reduced this impact.

The study did not consider the potential influence of pollutants or anthropogenic stressors on the morphology or growth of sugar kelp. However, since the study sites were far from any significant pollution sources and there were no reported pollution events in the local area during the study period, these factors were not expected to have influenced the results.

#### 4.1.8 Influence of kelp canopy and farm infrastructure on seabed shading

The discrepancy in shading between Carn a Wig in 2022 and St. Justinian in 2023 can be largely attributed to the differences in infrastructure used. At Carn a Wig, large 200L mussel buoys spaced just 7m apart were used in the long-line design, whereas St. Justinian used much smaller 13L buoys spaced 12m apart, resulting in considerably less shading, regardless of seaweed growth. It is also important to note that adjacent seaweed lines at St. Justinian were set at 15m lateral spacing, and therefore the shading directly below any cultivation line will receive minimal shading from adjacent lines. These findings suggest that the choice of infrastructure and farm design is a key driver for shading beneath the farms, with larger buoys and denser spacing of parallel lines contributing to more extensive shading and potentially influencing the undersea light-dependent ecological processes.

Previous studies on a sugar kelp farm found up to 40% decrease in light irradiance 5 meters below the seaweed farm where culture lines were spaced just 4m apart, potentially affecting communities beneath, particularly more sensitive habitats, such as seagrass meadows or maerl beds [2], [20]. Nevertheless, as the peak biomass period in a seaweed farming season is short and typically happens towards the end of the season, the ecological impact on photosynthetic benthic communities is likely minimal. However, it is wise to steer clear of densely vegetated areas or protected habitats to mitigate potential compounding effect on primary production, especially as the scale and density of seaweed farms increase [55].

#### 4.1.9 Influence of environmental parameters on biofouling

Biofouling has been identified previously as a key constraint for many seaweed farms worldwide. In this study, biofouling was significantly lower at Carn a Wig– a sheltered site with high tidal currents, compared to the adjacent site Porthlysgi– an exposed site with lower tidal currents. The degree of biofouling is typically associated with the relative wave exposure of locations, with more wave-exposed sites typically showing less biofouling than sheltered sites [20]. However, our study suggests that the correlation is not necessarily associated with wave-exposure, but rather that current-exposure is a key driver, which has also been suggested in previous studies in the northeast Atlantic [56]. As highlighted earlier, our findings demonstrate that consistent high current speeds as provided by tidal straits, such as the Ramsey Sound, which is adjacent to the Carn a Wig site, not only regulate temperatures, but also attenuate light levels through sedimentation during the summer period. This buffering effect on environmental factors reduces the proliferation of biofouling, thereby extending the harvest window and improving crop quality for seaweed farmers.

As the frequency and duration of marine heat waves are expected to increase in the future, the optimal timing for harvest will become increasingly difficult to predict, especially in sites where low summertime current speeds allow light and temperature levels to intensify. This could have a significant impact on the seaweed farming industry, as it would make it more difficult to plan and manage harvests. Although further research is needed to better understand the influencing factors and mitigation strategies that can impact biofouling (for example the effect of seeding density, local epibiont environments and seaweed farm design), by siting seaweed farms in locations with regular tidal currents many of the disadvantages associated with biofouling can be minimised.

#### 4.1.10 Conclusions on direct observations and environmental conditions

Our study demonstrates that local geographical and climatic differences, specifically between the two study sites, significantly impact seaweed growth and farm operations. Key factors like light availability, sea surface temperature, and water motion are vital in determining seaweed morphology and productivity, guiding the development of adaptive farming practices. Additionally, our research underscores the significance of strategic site selection and farm infrastructure design, including buoy size and line spacing, to manage biofouling effectively and optimise yields. These factors also play a pivotal role in controlling seabed shading, and whilst the present study showed minimal shading to the seabed, we caution in setting up dense farm infrastructure near light-sensitive benthic habitats, such as seagrass meadows or maerl beds, to prevent ecological disturbances.

### 4.2 BRUV

This study represents the first use of BRUV to investigate the size and abundance of mobile fauna in the vicinity of an integrated multi-trophic aquaculture (IMTA) seaweed and shellfish farm and builds upon previous BRUV studies in Wales in the proximity to natural kelp habitats [29], [30], [31]. The study identified a total of 15 fish and 6 invertebrate species, of which 9 fish and 1 invertebrate species are commercially important in the UK (marked with * in Table 2), all of which except dogfish (used as bait) and sand eel (used as fish meal) are commonly consumed as seafood. While the current data cannot categorically describe these habitats as fish ‘nurseries’ since the data doesn’t compare the findings with alternative habitats. However, the data does support that the benthic habitats beneath seaweed farms provide commercially important juvenile fish habitats that are comparable to those provided by wild kelp reefs in the same region [29].

#### 4.2.1 Limitations of BRUV

Whilst the study found that BRUVs were effective in sampling fish and invertebrate assemblages, it is likely that results are conservative due to reduced detection of juvenile flatfish (such as plaice and brill) and cryptic species (such as sea scorpion and worm pipefish). These species are less attracted to bait positioned above the seafloor, which results in decreased likelihood of their appearance in the video footage.

Although Nmax is the most commonly used metric for examining relative fish abundance with baited camera systems, some studies have shown that Nmax may not always be an accurate estimate of relative abundance as this can be influenced by the duration of the stay at the bait with species exhibiting long staying times having higher high Nmax values [29].

Whilst BRUVs allow for monitoring of motile fauna within sensitive habitats, the use of bait results in the attraction of specific species that are more sensitive to detecting the presence of the bait. In particular, it has been found that the presence of bait increases abundance and species richness of generalist carnivores but does not influence herbivorous fish [57]. The use of BRUV is therefore not dissimilar to other fish sampling techniques, as conventional invasive forms of fish sampling, such as netting, which too have a level of bias [58].

#### 4.2.2 Comparison of species detected between study sites

A significant pattern in our data is that of differing faunal assemblages between the two sites. The presence of greater spotted dogfish (*S. stellaris*) was found to be unique to Porthlysgi where the substrate is mostly sand and occasional boulders with seaweed assemblages. Dogfish have been found to prefer sandy habitats due to the abundance of their preferred prey and the provision of suitable environment for reproduction and nursery grounds [59]. Pollack (*P. pollachius*) were a unique species in Carn a Wig. Pollock typically inhabit environments where temperatures do not exceed much beyond 12°C, with strong tidal currents and prefer habitats with low light levels such as deep water [60].

#### 4.2.3 Species associated with kelp habitats

From the species observed, the two species that are most closely associated with kelp habitats are likely to be the Ballan wrasse (*Labrus bergylta*) and the Shore crab (*Carcinus maenas*). Ballan wrasse are known to be associated with kelp forests, where they use the dense kelp blades as a refuge from predators and as a hunting ground for prey [61]. They are also known to feed on the invertebrates that live in and around the kelp, such as crabs and snails. In addition, they can help to control the abundance of other species in the ecosystem, including sea urchins, which can have a negative impact on kelp forests if their populations become too large [62].

Shore crabs are also commonly found in kelp habitats, where they feed on a variety of small invertebrates, including barnacles and small crabs, which are abundant in and around the kelp blades [63]. They are also known to use the kelp blades as a refuge from predators and as a hunting ground for prey. In addition, they are important prey for many other organisms in the ecosystem, including fish, birds, and marine mammals [64].

While some of the other species on the list may also be found in kelp habitats, the Ballan wrasse and Shore crab are most closely associated with these habitats due to their reliance on kelp for shelter, food, and survival. Although both species were detected frequently throughout the monitoring period, our data is not statistically significant to suggest a correlation between the presence of these species and the cultivation of kelps near the surface.

#### 4.2.4 BRUV conclusion

In conclusion, we find the benthic habitat below the seaweed farm areas to be important habitats for juvenile fish of commercial importance. Although all sampling locations had such a role, the extent of this was highly variable, with differing species assemblage patterns evident between sites. The present study provides evidence of the benthic motile fauna below two sea-farms in their trial stages and serves as a baseline from which we can assess the long-term impacts from sea-farming activity as operations intensify toward commercial scale.

### 4.3 PAM

#### 4.3.1 Comparison with previous PAM studies

When comparing to previous PAM studies, our measurements of DPM are comparable to that recorded in a location within 200m to our deployment site in July and August 2009 [65]. Although these studies used a similar PAM system their units were deployed inside fixed polyethylene pipes to allow POD’s to hang vertically in the high-current environment.

This arrangement may improve their detection probability and therefore it is possible that a higher DPM was recorded. Further, explanations for differences between these studies could also be down to differing intensities of boat traffic and varying hydrological complexities as further discussed in the following sections.

#### 4.3.2 Diurnal variation in cetacean activity

It is clear from our data that both harbour porpoises and bottlenose dolphins tended to visit and occupy certain locations near to the POD far more frequently at night than during the day, although the cause of this is unclear. A possible explanation may be that foraging intensifies at these locations at night, either because a higher density of prey was present or because there was more opportunity for prey capture (e.g. if prey emerged from refuges to feed at night). However, this pattern has not been observed universally. In a similar study in North Wales, a POD showed the opposite pattern [65]. Alternatively, porpoises may have avoided the study sites during the day, having been displaced for example, by high levels of boat disturbance. This idea is backed by the summer diurnal detection pattern, which reveals an almost total lack of detections during the day, corresponding with tourist boat activity in the region. In contrast, during winter, when there’s less tourism, the diurnal detection pattern shows a substantial number of daytime detections. Further study is required to determine whether prey availability is greater at these locations at night, and/or whether boat traffic is a contributory factor that explains the difference in levels of porpoise activity at some locations during the night and day.

#### 4.3.3 Tidal correlations of cetacean activity

The present study found no clear tidal correlations, despite previous studies in Ramsey Sound reporting regular patterns of activity correlating with tide state [34], [66]. A likely cause for this lack in correlation is due to the deployment site being in an area characterised by eddies, giving rise to a complex relationship between tide state and current speed. Indeed, previous studies have highlighted that porpoise spatiotemporal distribution variability is associated with hydrographic complexity features such as eddies, rips, upwellings, and drop-offs, which may concentrate prey and provide predictable foraging opportunities [67].

#### 4.4.4. Residency times

The study revealed that both harbour porpoises and bottlenose dolphins exhibited relatively short residency times near the seaweed farm (8-16 minutes), suggesting that both species were merely passing through the area rather than engaging in significant feeding activities.

The consistent residency times across different time periods (monthly and weekly) indicate a transient visitation pattern to the seaweed farm area, suggesting that the seaweed farm may not be a primary feeding site for these cetaceans or that other factors, such as boat traffic might influence their presence and activities in the area. Given the biodiversity-rich habitat of Ramsey Sound, it is expected that the relatively small expanse of the seaweed farm area, in comparison to the larger, rich feeding grounds available in Ramsey Sound, will have an insignificant impact on the foraging behaviour of cetaceans.

#### 4.4.5. Implications for monitoring and conservation

While POD data cannot estimate animal abundance, the high DPM for bottlenose dolphins indicates their abundance in Ramsey Sound. Previous PAM studies in the area predominantly reported harbour porpoises with only one report recording 4.6% dolphin detections [34], [68]. In a similar study approximately 15 miles along the coast from Ramsey Sound, near Fishguard, eight of ten POD locations showed that dolphins are more abundant in the summer and porpoises in winter [35]. Moreover, this same study showed that even when PODs are located only 1 km apart, completely different patterns of bottlenose dolphin occurrence can be detected. This calls for several PODs covering larger areas in future monitoring studies.

#### 4.4.6. PAM conclusions

In conclusion, we find a consistent presence harbour porpoise and a slightly higher presence of dolphins at the trial sea-farms compared with previous studies. We observed a strong difference between night-time and day-time presence of both species. However, to determine whether this variation is caused by natural drivers or anthropological impacts will require more detailed long-term POD measurement data capturing both cetaceans and boat sonar signals. These variations between night and day and the tidal-cycle can be used to help guide operational scheduling at the sea-farms to minimise disturbance of cetaceans. Given that there is currently no article or research, to our knowledge, that specifically investigates the correlation between cetaceans and their behaviour in relation to seaweed farms, this is an area that warrants further research. As seaweed farms scale further offshore, where there are fewer competing wild feeding habitats and the size of the farms are comparatively large, understanding the impacts on cetacean behaviour becomes increasingly important for conservation and management practices.

### 5. Conclusion

This study explored the environmental and biodiversity impacts of multiple small-scale seaweed farms in Pembrokeshire, UK, using affordable and user-friendly monitoring methods. It was observed that seaweed farming structures serve as stable substrates for biodiversity accumulation, with no negative impacts on marine biodiversity observed within the monitoring period, even as farming activities were ongoing. Despite the dynamic environmental conditions, small scale of operations, and short monitoring timeframe, the study sets a crucial baseline for future research, emphasising the need for routine, integrated biodiversity monitoring in seaweed farming operations. The findings underscore the potential of seaweed farms to support marine biodiversity, highlighting their role in sustainable marine management and the importance of further research to fully understand their ecological impacts, especially as seaweed farms expand into larger offshore areas. Moreover, the insights gained hold the potential to pave the way for resilient seaweed aquaculture industry in the face of climate change.

## Acknowledgements

PEBL CIC conducted the measurements in 2022 and 2023 at Car-Y-Mor sea-farms, and thanks O. Haines, F. Evans, S. Rees, M. Charlton for logistic support. This work was carried out as part of the ‘Car-Y-Mor Ecological Monitoring’ project, funded by WWF.

## References

[1] FAO., ‘The State of World Fisheries and Aquaculture 2020: Sustainability in Action; The State of World Fisheries and Aquaculture (SOFIA); FAO: Rome, Italy, 2020; ISBN 978-92-5-132692-3.’ 2020.

[2] I. Campbell et al., ‘The Environmental Risks Associated With the Development of Seaweed Farming in Europe - Prioritizing Key Knowledge Gaps’, Front. Mar. Sci., vol. 6, p. 107, Mar. 2019, doi: 10.3389/fmars.2019.00107.

[3] H. Forbes, V. Shelamoff, W. Visch, and C. Layton, ‘Farms and forests: evaluating the biodiversity benefits of kelp aquaculture’, J Appl Phycol, vol. 34, no. 6, pp. 3059–3067, Dec. 2022, doi: 10.1007/s10811-022-02822-y.

[4] L. Hasselström, W. Visch, F. Gröndahl, G. M. Nylund, and H. Pavia, ‘The impact of seaweed cultivation on ecosystem services - a case study from the west coast of Sweden’, Marine Pollution Bulletin, vol. 133, pp. 53–64, Aug. 2018, doi: 10.1016/j.marpolbul.2018.05.005.

5. R. Langton, S. Augyte, N. Price, J. Forster, T. Noji, and G. Grebe, ‘An Ecosystem Approach to the Culture of Seaweed’, 2019.

[6] M. Y. Roleda and C. L. Hurd, ‘Seaweed nutrient physiology: application of concepts to aquaculture and bioremediation’, Phycologia, vol. 58, no. 5, pp. 552–562, Sep. 2019, doi: 10.1080/00318884.2019.1622920.

[7] F. W. R. Ross et al., ‘Potential role of seaweeds in climate change mitigation’, Science of The Total Environment, vol. 885, p. 163699, Aug. 2023, doi: 10.1016/j.scitotenv.2023.163699.

[8] S. Spillias et al., ‘The empirical evidence for the social-ecological impacts of seaweed farming’, PLOS Sustain Transform, vol. 2, no. 2, p. e0000042, Feb. 2023, doi: 10.1371/journal.pstr.0000042.

9. [9] DEFRA, ‘Consultation on the Principles of Marine Net Gain’. [Online]. Available: https://consult.defra.gov.uk/defra-net-gain-consultation-team/consultation-on-the-principles-of-marine-net-gain/

[10] A. McQuatters-Gollop et al., ‘Assessing the state of marine biodiversity in the Northeast Atlantic’, Ecological Indicators, vol. 141, p. 109148, Aug. 2022, doi: 10.1016/j.ecolind.2022.109148.

[11] C. M. Duarte, A. Bruhn, and D. Krause-Jensen, ‘A seaweed aquaculture imperative to meet global sustainability targets’, Nat Sustain, vol. 5, no. 3, pp. 185–193, Oct. 2021, doi: 10.1038/s41893-021-00773-9.

[12] D. S. Schmeller et al., ‘Building capacity in biodiversity monitoring at the global scale’, Biodivers Conserv, vol. 26, no. 12, pp. 2765–2790, Nov. 2017, doi: 10.1007/s10531-017-1388-7.

[13] I. Fitridge, T. Dempster, J. Guenther, and R. De Nys, ‘The impact and control of biofouling in marine aquaculture: a review’, Biofouling, vol. 28, no. 7, pp. 649–669, Aug. 2012, doi: 10.1080/08927014.2012.700478.

[14] J. Bannister, M. Sievers, F. Bush, and N. Bloecher, ‘Biofouling in marine aquaculture: a review of recent research and developments’, Biofouling, vol. 35, no. 6, pp. 631–648, Jul. 2019, doi: 10.1080/08927014.2019.1640214.

[15] C. S. Park and E. K. Hwang, ‘Seasonality of epiphytic development of the hydroid Obelia geniculata on cultivated Saccharina japonica (Laminariaceae, Phaeophyta) in Korea’, J Appl Phycol, vol. 24, no. 3, pp. 433–439, Jun. 2012, doi: 10.1007/s10811-011-9755-3.

[16] C. L. Hurd, K. M. Durante, F.-S. Chia, and P. J. Harrison, ‘Effect of bryozoan colonization on inorganic nitrogen acquisition by the kelps Agarum fimbriatum and Macrocystis integrifolia’, Marine Biology, vol. 121, no. 1, pp. 167–173, Dec. 1994, doi: 10.1007/BF00349486.

[17] D. Pica et al., ‘Dynamics of a biofouling community in finfish aquaculture: a case study from the South Adriatic Sea’, Biofouling, vol. 35, no. 6, pp. 696–709, Jul. 2019, doi: 10.1080/08927014.2019.1652817.

[18] T. Bekkby et al., ‘“Hanging gardens”—comparing fauna communities in kelp farms and wild kelp forests’, Front. Mar. Sci., vol. 10, p. 1066101, Feb. 2023, doi: 10.3389/fmars.2023.1066101.

[19] A. Walls, R. Kennedy, R. Fitzgerald, A. Blight, M. Johnson, and M. Edwards, ‘Potential novel habitat created by holdfasts from cultivated Laminaria digitata: assessing the macroinvertebrate assemblages’, Aquacult. Environ. Interact., vol. 8, pp. 157–169, Feb. 2016, doi: 10.3354/aei00170.

[20] W. Visch, M. Kononets, P. O. J. Hall, G. M. Nylund, and H. Pavia, ‘Environmental impact of kelp (Saccharina latissima) aquaculture’, Marine Pollution Bulletin, vol. 155, p. 110962, Jun. 2020, doi: 10.1016/j.marpolbul.2020.110962.

[21] H. Christie, G. S. Andersen, L. A. Tveiten, and F. E. Moy, ‘Macrophytes as habitat for fish’, ICES Journal of Marine Science, vol. 79, no. 2, pp. 435–444, Mar. 2022, doi: 10.1093/icesjms/fsac008.

[22] M. W. Beck et al., ‘The Identification, Conservation, and Management of Estuarine and Marine Nurseries for Fish and Invertebrates’, BioScience, vol. 51, no. 8, p. 633, 2001, doi: 10.1641/0006-3568(2001)051[0633:TICAMO]2.0.CO;2.

[23] S. Corrigan et al., ‘Home sweet home: Comparison of epibiont assemblages associated with cultivated and wild sugar kelp (Saccharina latissima), co-cultivated blue mussels (Mytilus edulis) and farm infrastructure’, In Review, preprint, May 2023. doi: 10.21203/rs.3.rs-2934251/v1.

[24] C. Chen, T. A. Jefferson, B. Chen, and Y. Wang, ‘Geographic range size, water temperature, and extrinsic threats predict the extinction risk in global cetaceans’, Global Change Biology, vol. 28, no. 22, pp. 6541–6555, Nov. 2022, doi: 10.1111/gcb.16385.

[25] A. Azzellino et al., ‘An index based on the biodiversity of cetacean species to assess the environmental status of marine ecosystems’, Marine Environmental Research, vol. 100, pp. 94– 111, Sep. 2014, doi: 10.1016/j.marenvres.2014.06.003.

[26] E. Parsons et al., ‘Key research questions of global importance for cetacean conservation’, Endang. Species. Res., vol. 27, no. 2, pp. 113–118, Feb. 2015, doi: 10.3354/esr00655.

27. [27] Steven Hermans, ‘2023 Seaweed State of the Industry’. [Online]. Available: https://phyconomy.net/articles/2022-seaweed-review/

[28] G. E. Bath, C. A. Price, K. L. Riley, and J. A. Morris, ‘A global review of protected species interactions with marine aquaculture’, Reviews in Aquaculture, vol. 15, no. 4, pp. 1686–1719, Sep. 2023, doi: 10.1111/raq.12811.

[29] R. K. F. Unsworth, J. R. Peters, R. M. McCloskey, and S. L. Hinder, ‘Optimising stereo baited underwater video for sampling fish and invertebrates in temperate coastal habitats’, Estuarine, Coastal and Shelf Science, vol. 150, pp. 281–287, Oct. 2014, doi: 10.1016/j.ecss.2014.03.020.

[30] R. E. Jones, R. A. Griffin, S. C. Rees, and R. K. F. Unsworth, ‘Improving visual biodiversity assessments of motile fauna in turbid aquatic environments’, Limnology & Ocean Methods, vol. 17, no. 10, pp. 544–554, Oct. 2019, doi: 10.1002/lom3.10331.

[31] R. E. Jones, R. A. Griffin, R. J. H. Herbert, and R. K. F. Unsworth, ‘Consistency Is Critical for the Effective Use of Baited Remote Video’, Oceans, vol. 2, no. 1, pp. 215–232, Mar. 2021, doi: 10.3390/oceans2010013.

[32] E. Furness and R. K. F. Unsworth, ‘Demersal Fish Assemblages in NE Atlantic Seagrass and Kelp’, Diversity, vol. 12, no. 10, p. 366, Sep. 2020, doi: 10.3390/d12100366.

[33] P. G. H. Evans and P. S. Hammond, ‘Monitoring cetaceans in European waters’, Mammal Review, vol. 34, no. 1–2, pp. 131–156, Jan. 2004, doi: 10.1046/j.0305-1838.2003.00027.x.

[34] C. Pierpoint, ‘Harbour porpoise (*Phocoena phocoena*) foraging strategy at a high energy, near-shore site in south-west Wales, UK’, J. Mar. Biol. Ass., vol. 88, no. 6, pp. 1167–1173, Sep. 2008, doi: 10.1017/S0025315408000507.

[35] M. Simon, H. Nuuttila, M. M. Reyes-Zamudio, F. Ugarte, U. Verfub, and P. G. H. Evans, ‘Passive acoustic monitoring of bottlenose dolphin and harbour porpoise, in Cardigan Bay, Wales, with implications for habitat use and partitioning’, J. Mar. Biol. Ass., vol. 90, no. 8, pp. 1539–1545, Dec. 2010, doi: 10.1017/S0025315409991226.

36. J. D. J. Macaulay, ‘Passive Acoustic Monitoring of Harbour Porpoise Behaviour, Distribution and Density in Tidal Rapid Habitats’.

[37] H. K. Nuuttila, W. Courtene-Jones, S. Baulch, M. Simon, and P. G. H. Evans, ‘Don’t forget the porpoise: acoustic monitoring reveals fine scale temporal variation between bottlenose dolphin and harbour porpoise in Cardigan Bay SAC’, Mar Biol, vol. 164, no. 3, p. 50, Mar. 2017, doi: 10.1007/s00227-017-3081-5.

[38] L. A. Kyhn et al., ‘From echolocation clicks to animal density—Acoustic sampling of harbor porpoises with static dataloggers’, The Journal of the Acoustical Society of America, vol. 131, no. 1, pp. 550–560, Jan. 2012, doi: 10.1121/1.3662070.

[39] D. A. Smale et al., ‘Marine heatwaves threaten global biodiversity and the provision of ecosystem services’, Nat. Clim. Chang., vol. 9, no. 4, pp. 306–312, Apr. 2019, doi: 10.1038/s41558-019-0412-1.

[40] S. Corrigan et al., ‘Development and Diversity of Epibiont Assemblages on Cultivated Sugar Kelp (Saccharina latissima) in Relation to Farming Schedules and Harvesting Techniques’, Life, vol. 13, no. 1, p. 209, Jan. 2023, doi: 10.3390/life13010209.

[41] L. A. Green and C. D. Neefus, ‘Effects of temperature, light level, photoperiod, and ammonium concentration on Pyropia leucosticta (Bangiales, Rhodophyta) from the Northwest Atlantic’, J Appl Phycol, vol. 27, no. 3, pp. 1253–1261, Jun. 2015, doi: 10.1007/s10811-014-0421-4.

[42] K. Bischof et al., ‘Kelps and Environmental Changes in Kongsfjorden: Stress Perception and Responses’, in The Ecosystem of Kongsfjorden, Svalbard, vol. 2, H. Hop and C. Wiencke, Eds., in Advances in Polar Ecology, vol. 2., Cham: Springer International Publishing, 2019, pp. 373–422. doi: 10.1007/978-3-319-46425-1_10.

[43] J. Nepper-Davidsen, D. Andersen, and M. Pedersen, ‘Exposure to simulated heatwave scenarios causes long-term reductions in performance in Saccharina latissima’, Mar. Ecol. Prog. Ser., vol. 630, pp. 25–39, Nov. 2019, doi: 10.3354/meps13133.

[44] I. R. Davison, ‘Adaptation of photosynthesis in Laminaria Saccharina to changes in growth temperature’, Journal of Phycology, vol. 23, no. s2, pp. 273–283, Jun. 1987, doi: 10.1111/j.1529-8817.1987.tb04135.x.

[45] N. Diehl, M. Y. Roleda, I. Bartsch, U. Karsten, and K. Bischof, ‘Summer Heatwave Impacts on the European Kelp Saccharina latissima Across Its Latitudinal Distribution Gradient’, Front. Mar. Sci., vol. 8, p. 695821, Oct. 2021, doi: 10.3389/fmars.2021.695821.

[46] F. Goecke, G. Klemetsdal, and Å. Ergon, ‘Cultivar Development of Kelps for Commercial Cultivation—Past Lessons and Future Prospects’, Front. Mar. Sci., vol. 8, p. 110, Feb. 2020, doi: 10.3389/fmars.2020.00110.

[47] G. S. Grebe, C. J. Byron, D. C. Brady, A. T. St. Gelais, and B.A. Costa-Pierce, ‘The effect of distal-end trimming on Saccharina latissima morphology, composition, and productivity’, J World Aquaculture Soc, vol. 52, no. 5, pp. 1081–1098, Oct. 2021, doi: 10.1111/jwas.12814.

[48] L. Kregting, A. J. Blight, B. Elsäßer, and G. Savidge, ‘The influence of water motion on the growth rate of the kelp Laminaria digitata’, Journal of Experimental Marine Biology and Ecology, vol. 478, pp. 86–95, May 2016, doi: 10.1016/j.jembe.2016.02.006.

[49] J. C. Sanderson, M. J. Dring, K. Davidson, and M. S. Kelly, ‘Culture, yield and bioremediation potential of Palmaria palmata (Linnaeus) Weber & Mohr and Saccharina latissima (Linnaeus) C.E. Lane, C. Mayes, Druehl & G.W. Saunders adjacent to fish farm cages in northwest Scotland’, Aquaculture, vol. 354–355, pp. 128–135, Jul. 2012, doi: 10.1016/j.aquaculture.2012.03.019.

[50] M. A. R. Koehl, W. K. Silk, H. Liang, and L. Mahadevan, ‘How kelp produce blade shapes suited to different flow regimes: A new wrinkle’, Integrative and Comparative Biology, vol. 48, no. 6, pp. 834–851, Apr. 2008, doi: 10.1093/icb/icn069.

[51] T. Bekkby, E. Rinde, H. Gundersen, K. Norderhaug, J. Gitmark, and H. Christie, ‘Length, strength and water flow: relative importance of wave and current exposure on morphology in kelp Laminaria hyperborea’, Mar. Ecol. Prog. Ser., vol. 506, pp. 61–70, Jun. 2014, doi: 10.3354/meps10778.

[52] S. Matsson, A. Metaxas, S. Forbord, S. Kristiansen, A. Handå, and B. A. Bluhm, ‘Effects of outplanting time on growth, shedding and quality of Saccharina latissima (Phaeophyceae) in its northern distribution range’, J Appl Phycol, vol. 33, no. 4, pp. 2415–2431, Aug. 2021, doi: 10.1007/s10811-021-02441-z.

[53] A. H. Buschmann et al., ‘Seaweed production: overview of the global state of exploitation, farming and emerging research activity’, European Journal of Phycology, vol. 52, no. 4, pp. 391–406, Oct. 2017, doi: 10.1080/09670262.2017.1365175.

[54] P. D. Kerrison et al., ‘Twine selection is essential for successful hatchery cultivation of Saccharina latissima, seeded with either meiospores or juvenile sporophytes’, J Appl Phycol, vol. 31, no. 5, pp. 3051–3060, Oct. 2019, doi: 10.1007/s10811-019-01793-x.

[55] E. J. Cottier-Cook et al., ‘A new Progressive Management Pathway for improving seaweed biosecurity’, Nat Commun, vol. 13, no. 1, p. 7401, Dec. 2022, doi: 10.1038/s41467-022-34783-8.

[56] A. Mols-Mortensen, E. Á. G. Ortind, C. Jacobsen, and S. L. Holdt, ‘Variation in growth, yield and protein concentration in Saccharina latissima (Laminariales, Phaeophyceae) cultivated with different wave and current exposures in the Faroe Islands’, J Appl Phycol, vol. 29, no. 5, pp. 2277–2286, Oct. 2017, doi: 10.1007/s10811-017-1169-4.

[57] T. Langlois, E. Harvey, B. Fitzpatrick, J. Meeuwig, G. Shedrawi, and D. Watson, ‘Cost-efficient sampling of fish assemblages: comparison of baited video stations and diver video transects’, *Aquat*. Biol., vol. 9, no. 2, pp. 155–168, Apr. 2010, doi: 10.3354/ab00235.

[58] G. English et al., ‘A review of data collection methods used to monitor the associations of wild species with marine aquaculture sites’, *Reviews in Aquaculture*, p. raq.12890, Jan. 2024, doi: 10.1111/raq.12890.

[59] J. R. Ellis, S. R. McCully Phillips, and F. Poisson, ‘A review of capture and post-release mortality of elasmobranchs’, Journal of Fish Biology, vol. 90, no. 3, pp. 653–722, Mar. 2017, doi: 10.1111/jfb.13197.

[60] J. H. Ólafsdóttir, J. G. Þorbjörnsson, B. K. Kristjánsson, and J. S. Ólafsson, ‘Invertebrate biodiversity in cold groundwater fissures in Iceland’, Ecology and Evolution, vol. 9, no. 11, pp. 6399–6409, Jun. 2019, doi: 10.1002/ece3.5213.

[61] K. M. Norderhaug, H. Christie, J. H. Fosså, and S. Fredriksen, ‘fish–macrofauna interactions in a kelp (*laminaria hyperborea*) forest’, J. Mar. Biol. Ass., vol. 85, no. 5, pp. 1279–1286, Oct. 2005, doi: 10.1017/S0025315405012439.

[62] S. J. Bourlat et al., ‘Wrasse fishery on the Swedish West Coast: towards ecosystem-based management’, ICES Journal of Marine Science, vol. 78, no. 4, pp. 1386–1397, Aug. 2021, doi: 10.1093/icesjms/fsaa249.

[63] P. A. Todd, R. A. Briers, R. J. Ladle, and F. Middleton, ‘Phenotype-environment matching in the shore crab (Carcinus maenas)’, Marine Biology, vol. 148, no. 6, pp. 1357–1367, Apr. 2006, doi: 10.1007/s00227-005-0159-2.

[64] F. Bulleri, ‘Role of recruitment in causing differences between intertidal assemblages on seawalls and rocky shores’, Mar. Ecol. Prog. Ser., vol. 287, pp. 53–65, 2005, doi: 10.3354/meps287053.

65. Welsh Assembly Government, ‘Assessment of Risk to Marine Mammals from Underwater Marine Renewable Devices in Welsh waters’. 2011.

[66] C. Malinka, D. Gillespie, J. Macaulay, R. Joy, and C. Sparling, ‘First in situ passive acoustic monitoring for marine mammals during operation of a tidal turbine in Ramsey Sound, Wales’, Mar. Ecol. Prog. Ser., vol. 590, pp. 247–266, Mar. 2018, doi: 10.3354/meps12467.

[67] F. M. Van Beest et al., ‘Environmental drivers of harbour porpoise fine-scale movements’, Mar Biol, vol. 165, no. 5, p. 95, May 2018, doi: 10.1007/s00227-018-3346-7.

[68] D. Gillespie, L. Palmer, J. Macaulay, C. Sparling, and G. Hastie, ‘Passive acoustic methods for tracking the 3D movements of small cetaceans around marine structures’, PLoS ONE, vol. 15, no. 5, p. e0229058, May 2020, doi: 10.1371/journal.pone.0229058.

